# Pglyrp1-Cre Marks Distinct Epithelial and Immune Lineages Across Mucosal Sites

**DOI:** 10.64898/2025.12.22.696025

**Authors:** Samuel Alvarez-Arguedas, Michael U. Shiloh

**Affiliations:** Department of Internal Medicine, University of Texas Southwestern Medical Center, Dallas, Texas 75390, USA; Department of Microbiology, University of Texas Southwestern Medical Center, Dallas, Texas 75390, USA

**Keywords:** Pglyrp1-Cre, M cells, Mucosal immunity

## Abstract

Mucosa-associated lymphoid tissue (MALT) initiates immune responses at mucosal entry sites. Within MALT, microfold (M) cells sample luminal antigens and deliver them to underlying immune cells. Despite their functional importance, few tools enable selective manipulation of M cells *in vivo*. Here we report the generation and characterization of a peptidoglycan recognition protein 1 (*Pglyrp1*)-Cre knock-in mouse designed to allow conditional genetic access to M cells. Using Rosa26-tdTomato reporter mice, we found strong *Pglyrp1* promoter activity in gut epithelial cells, including goblet and M cells, whereas activity in nasal-associated lymphoid tissue (NALT) was more heterogeneous and skewed towards immune cells, particularly neutrophils. To functionally interrogate *Pglyrp1*-expressing cells, we performed Cre-mediated ablation using three DTA-based models. The Rosa26^GFP-DTA^ line caused marked perinatal lethality in double-positive pups, suggesting essential roles for Pglyrp1-positive cells early in life. In contrast, Rosa26^DTA^ and Rosa26^iDTR^ crosses produced minimal depletion of mucosal populations, including M cells, even at the highest non-lethal diphtheria toxin dose. These findings demonstrate tissue-specific *Pglyrp1* promoter activity and highlight challenges in achieving M cell-specific targeting. Although not M cell-restricted, the Pglyrp1-Cre mouse provides a useful tool for manipulating Pglyrp1-expressing lineages and probing their roles in mucosal homeostasis and immunity.

## Introduction

Mucosa-associated lymphoid tissue (MALT) plays a central role in immune defense at major sites of pathogen entry, including the gastrointestinal (GI) tract, respiratory system, skin, and genitourinary tract [1]. MALT consist of organized lymphoid follicles covered by specialized epithelial layers, with Peyer’s patches serving as the best-characterized example [2]. From studies of Peyer’s patches, microfold (M) cells were identified as rare epithelial cells that sample luminal antigens and deliver them to subepithelial immune cells through transcytosis [3]. This process is exploited by pathogens such as *Salmonella, Yersinia, Shigella*, and *Mycobacterium tuberculosis* to enter the host [4–7]. M cells therefore play critical roles in microbial pathogenesis and mucosal immune surveillance [8].

Despite their importance, tools that selectively target M cells *in vivo* remain limited. Approaches have included disrupting RANK-RANKL signaling to modulate M cell abundance [9], or using mucosal adjuvants such as cholera toxin [10]. Genetically engineered models such as *Spib*- or *Sox8*-deficient mice [11, 12] and *Villin*-Cre-driven conditional knockouts targeting *Tnfrsf11a* (RANK), Gp2, or Atoh8 [13–16] have advanced the field, yet many lack cell-type specificity and affect broader epithelial compartments.

Peptidoglycan recognition protein 1 (PGLYRP1), also known as PGRP, PGRP-S, TAG7, or TNFAF3L, is an innate immune receptor expressed in several cell types. PGLYRP1 has been detected in M cells of Peyer’s patches, where it colocalizes with M cell markers such as *Ulex europaeus* agglutinin 1 (UEA-1) and GP2 [10, 17], and in RANKL-induced M cells derived from intestinal organoids [18]. PGLYRP1 is also expressed in neutrophils, epithelial cells, and stromal populations [19–21]. These observations suggested *Pglyrp1* as a potential promoter for M cell-specific Cre expression.

Here we describe the generation and characterization of a *Pglyrp1*-Cre knock-in mouse. Although initially intended for M cell targeting, *Pglyrp1*-Cre drove recombination in a broader set of epithelial and immune populations in a tissue-specific manner. Using fluorescent reporter and diphtheria toxin-based ablation models, we mapped *Pglyrp1-*expressing lineages and uncovered their functional significance. While not M cell-specific, this model provides a versatile tool for manipulating *Pglyrp1*-positive cells *in vivo* and for probing their contributions to mucosal barrier integrity and innate immunity.

## Results

### Generation of a *Pglyrp1*-Cre mouse model and founder screening

To genetically target *Pglyrp1*-expressing cells, we generated a knock-in mouse in which Cre recombinase was inserted downstream of exon 3 of the *Pglyrp1* locus using the Easi-CRISPR method [22]. This approach preserves endogenous promoter regulation while minimizing disruption to *Pglyrp1* gene function (Figure 1A). The single-stranded DNA (ssDNA) donor used for recombination included homology arms, an internal ribosome entry site (IRES), and the Cre coding sequence (Figure 1B). The full recombined exon 3 sequence and donor templates are shown in Supplemental Figures 1A and B.

**Figure 1.**
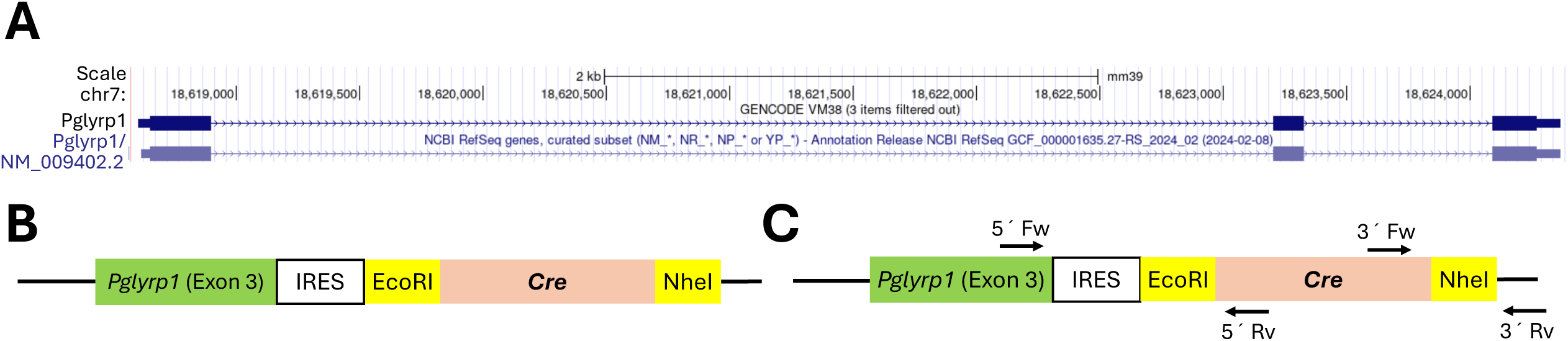
Generation of a *Pglyrp1*-Cre mouse model and founder screening. (A) Genomic sequence of mouse chromosome 7 showing the *Pglyrp1* gene, as visualized in the UCSC Genome browser. (B) Schematic representation of the 2,015bp ssDNA donor construct, illustrating all elements included for the targeted knock-in. (C) Primer locations used to identify the Cre recombinase insertion at the 5⍰ and 3⍰ ends of the modified genomic sequence. IRES = Internal ribosome entry site.

Founder animals were born at expected Mendelian ratios, were phenotypically normal, and remained fertile. For genotyping we used primer pairs spanning the 5⍰ and 3⍰ junctions of the insertion (Figure 1C) that generated distinct PCR patterns for wild-type and knock-in alleles. Restriction enzyme digestion with EcoRI and NheI further confirmed correct cassette integration (Supplemental Figures 1C-1E).

These analyses verified successful generation of the *Pglyrp1*-Cre knock-in line.

### Reporter expression in newborn *Pglyrp1*-Cre mice reveals tissue-specific patterns

To examine *Pglyrp1*-driven Cre activity, we crossed *Pglyrp1*-Cre mice with B6.Cg-Gt(ROSA)26Sor^tm14(CAG-tdTomato)Hze^/J (also known as Rosa26-tdTomato) reporter mice [23]. To examine whole-body reporter expression during early development, one-day-old pups were fixed, sectioned, and stained for RFP and the M cell marker GP2 [17].

The GI tract exhibited strong Tomato expression in *Pglyrp1*^Cre/wt^Rosa26^tdTom/wt^ pups, with clear colocalization between Tomato and GP2 (Figure 2A; Supplemental Figure 2A). Control mice lacking Cre showed only GP2 staining. A closer examination of the GI tract showed that Tomato-only controls lacked overlap between Tomato and GP2 signals, with only GP2-positive cells visible (Figure 2A, top row, open arrowheads). In contrast, colocalization of Tomato and GP2 (asterisks) was observed exclusively in the presence of Cre recombinase (Figure 2A, bottom row). Notably, Tomato^+^GP2^−^ cells (closed arrowheads) were more abundant than Tomato^−^GP2^+^ cells (open arrowheads), indicating *Pglyrp1* expression in epithelial lineages beyond classical M cells.

**Figure 2.**
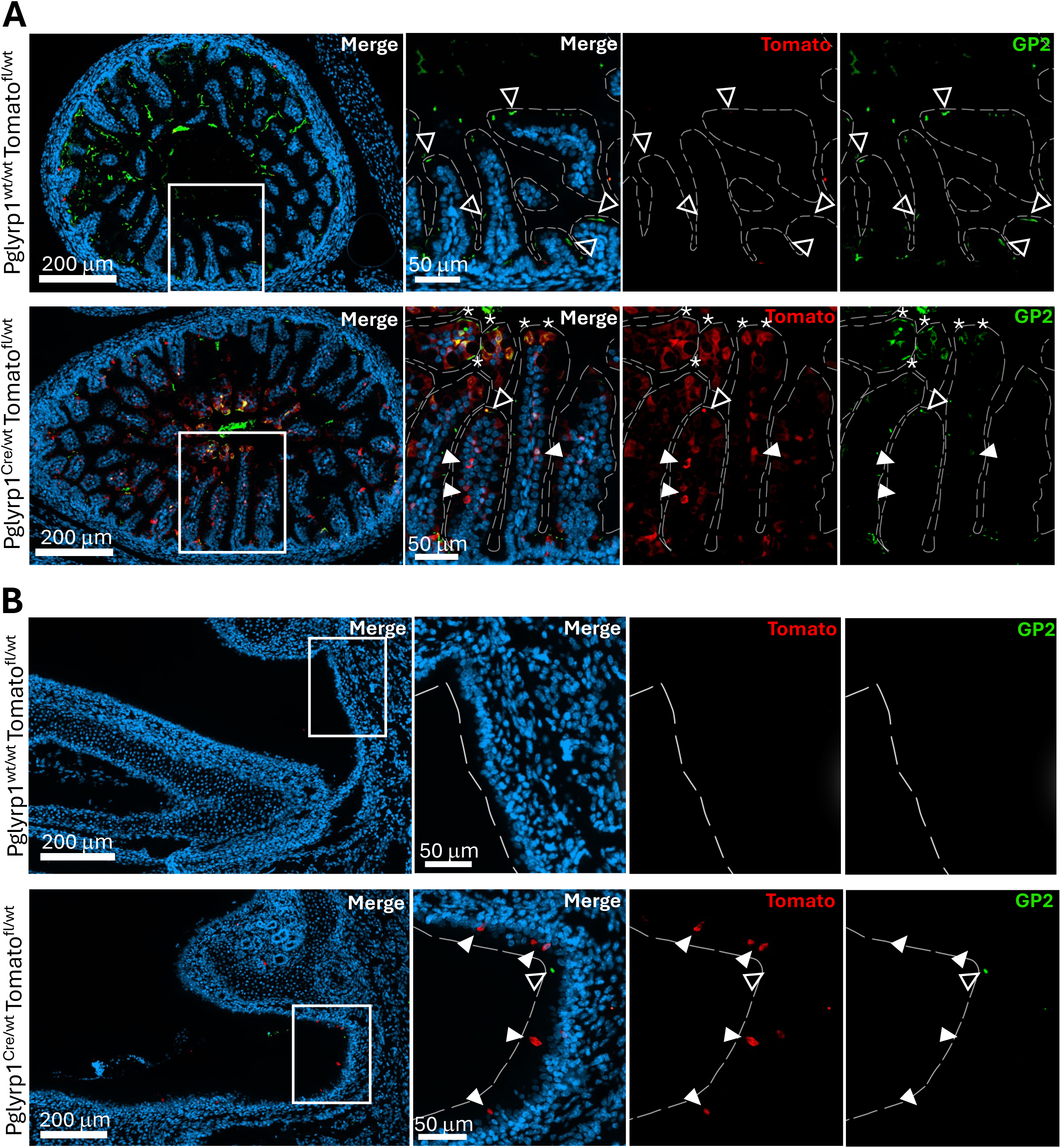
Reporter expression in newborn *Pglyrp1*-Cre mice reveals tissue-specific patterns. (A) Confocal images of representative Pglyrp1^wt/wt^ Rosa26^tdTom/wt^ (top), and Pglyrp1^Cre/wt^ Rosa26^tdTom/wt^ (bottom) one-day-old pups from the GI tissue (A), and NALT (B) sections stained with anti-red fluorescent protein (RFP), and anti-GP2 antibodies. The white box highlights the area shown at higher magnification in the right panels. Closed arrowhead points to Tomato^+^ GP2^-^ cells, open arrowhead to a Tomato^-^ GP2^+^ cells, and asterisk to Tomato^+^ GP2^+^ cells. Scale bars included in the pictures.

In the NALT, Tomato expression was detected only in Cre-positive mice (Figure 2B, bottom row, closed arrowheads). GP2 staining was rarely observed in either control mice or *Pglyrp1*-Cre mice, suggesting either very low M cell abundance at this stage (Figure 2B, Supplemental Figure 2B) or that GP2 is not a reliable marker for NALT M cells, consistent with recent findings in human airway tissue [24].

These observations indicate tissue-specific *Pglyrp1* promoter activity, with robust epithelial expression in the GI tract but limited or undetectable expression of canonical M cell markers such as GP2 in the upper airway mucosa.

### Quantitative analysis of *Pglyrp1*-Cre–driven Tomato expression in adult mice highlights tissue-specific patterns

To characterize *Pglyrp1*-driven reporter expression at single-cell resolution, we performed flow cytometry on GI tract and NALT tissues from adult mice (8-12 weeks of age) carrying Cre, the Tomato reporter, or both alleles. The antibody panel included markers for epithelial lineages, immune subsets, and M cell populations (Supplemental Figure 3A, orange boxes). Immune populations were included based on previous reports implicating PGLYRP1 in both innate and adaptive immunity, as well as its involvement in inflammatory responses [19–21, 25, 26].

Initial analyses confirmed that the presence of Cre recombinase allele, the Tomato reporter allele, or both did not alter the overall composition of major cell populations in either the GI tract (Supplemental Figure 3B) or the NALT (Supplemental Figure 3C). We detected Tomato fluorescence only in *Pglyrp1*^Cre/wt^ Rosa26td^Tom/wt^ animals, with strong expression across GI epithelial subsets (Figure 3A). Goblet cells (Cd45^-^Epcam^+^Gp2^+^Tnfaip2^-^), immature M cells (Cd45^-^Epcam^+^Gp2^-^ Tnfaip2^+^), and mature M cells (Cd45^-^Epcam^+^Gp2^+^Tnfaip2^+^) all exhibited very high Tomato positivity, indicating broad *Pglyrp1* promoter activity in gut epithelial lineages (Figure 3A).

**Figure 3.**
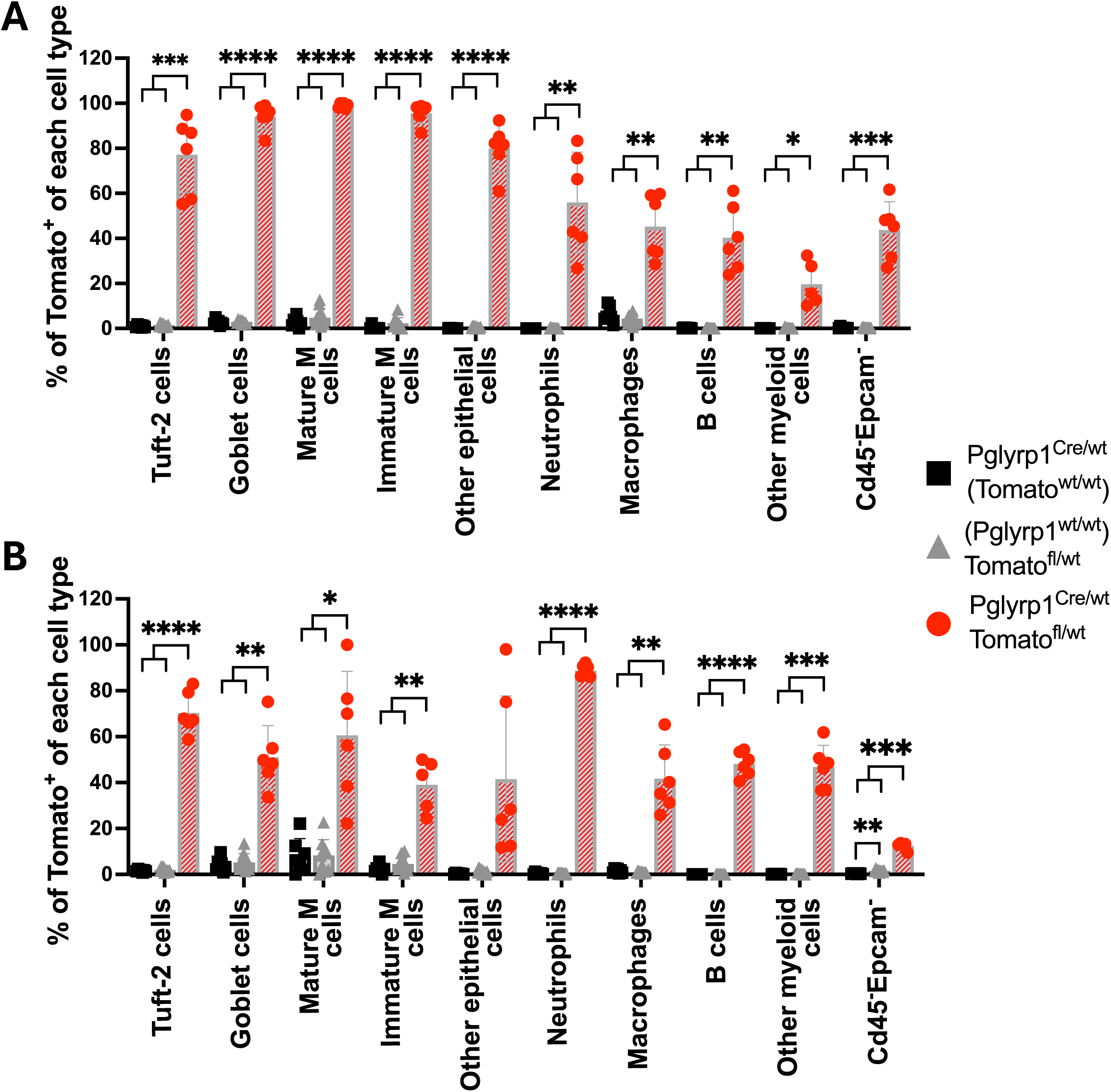
Quantitative analysis of Pglyrp1-Cre driven Tomato expression in adult mice highlights tissue-specific patterns. Flow cytometry analysis of Pglyrp1^Cre/wt^ (black squares), Rosa26^tdTom/wt^ (gray triangles), and Pglyrp1^Cre/wt^Rosa26^tdTom/wt^ (red circles) gut (A) and NALT (B) tissues in adult mice. Markers used to define each cell population are specified in the x-axis. Individual data are plotted with means ± SD analyzed with two-way ANOVA with Geisser-Greenhouse correction (no sphericity assumed). *p<0.05, **p<0.005, ***p<0.0005, ****p<0.0001. Where not shown, comparisons were not significant. N = 7 (Pglyrp1^Cre/wt^) and 10 (Rosa26^tdTom/wt^, Pglyrp1^Cre/wt^ Rosa26^tdTom/wt^). Markers used to identify each cell population are described in Supplemental Table 1A.

In contrast, Tomato expression in the NALT of Pglryp1^Cre/wt^Rosa26^tdTom/wt^ adult mice was more heterogeneous (Figure 3B). Neutrophils (Cd45^+^Epcam^-^Ly6g^+^) showed the highest reporter expression, while epithelial subsets displayed variable Tomato levels. Tuft cells (Cd45^+^Epcam^+^) showed the strongest epithelial expression (∼70%), whereas goblet cells (Cd45-Epcam+Gp2+Tnfaip2-) and M cell subsets, both immature (Cd45-Epcam+Gp2-Tnfaip2+) and mature (Cd45-Epcam+Gp2+Tnfaip2+), displayed intermediate expression (∼50%).

Together, these data reveal distinct tissue-specific patterns of *Pglyrp1* promoter activity: predominantly epithelial in the gut, and more immune-enriched in the NALT.

### *Pglyrp1*-Cre–Rosa26^DTA^ mice show no detectable changes in epithelial or immune composition

To test whether *Pglyrp1*-expressing populations could be depleted by constitutive diphtheria toxin expression, we crossed *Pglyrp1*-Cre mice with B6.129P2-Gt(ROSA)26Sor^tm1(DTA)Lky^/J (hereafter Rosa26^DTA^) mice, expected to drive cell-intrinsic diphtheria toxin (DTA) expression in Cre-positive lineages [27]. We analyzed the impact of this constitutive ablation approach on mucosal cell populations in the gut and NALT by flow cytometry, using a refined antibody panel designed to identify epithelial subsets, including GP2^+^ and GP2^−^ populations, and major immune lineages (Supplemental Figure 4A, orange boxes). Analysis by flow cytometry demonstrated no significant differences in epithelial or immune populations in GI tract or NALT across genotypes (Figure 4A, B). Neither GP2^−^ nor GP2^+^ epithelial populations were reduced.

**Figure 4.**
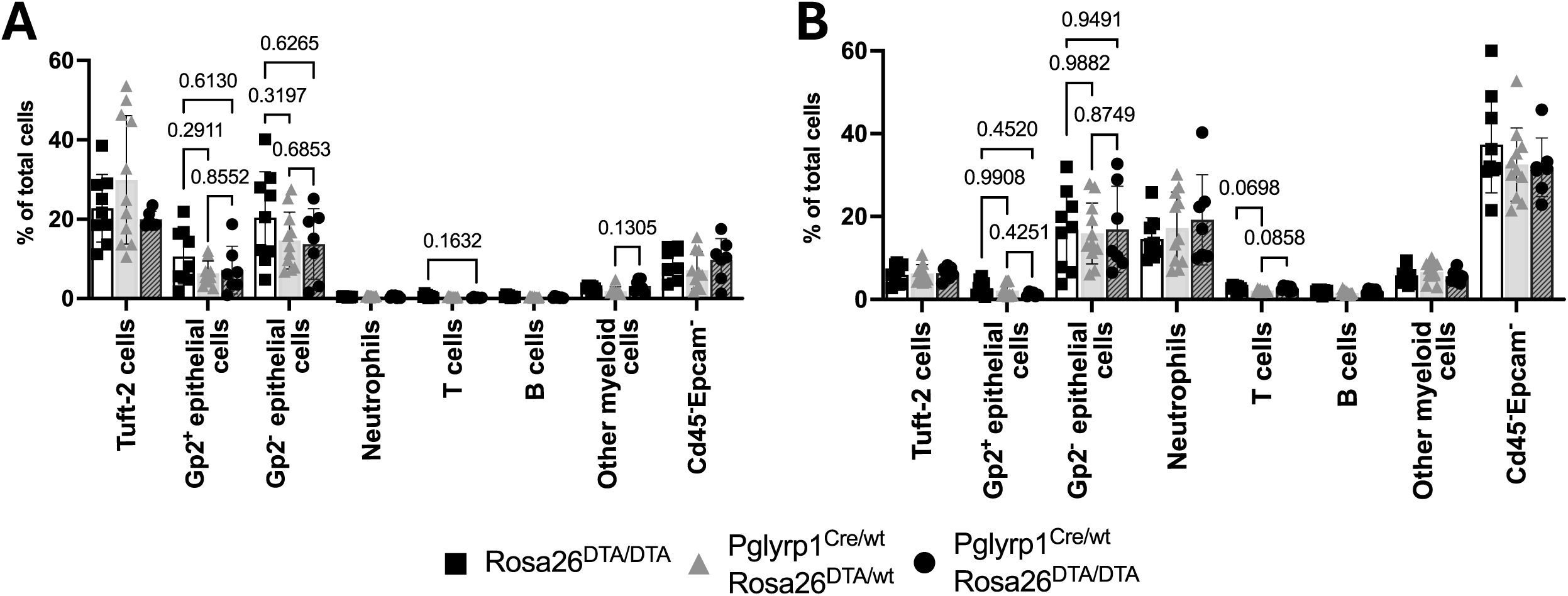
Pglyrp1-Cre–Rosa26^DTA^ mice show no detectable changes in epithelial or immune cell composition. Percentage of total cells in gut (A), and NALT (B) in Rosa26^DTA/DTA^ (black squares), Pglyrp1^Cre/wt^ Rosa26^DTA/wt^ (gray triangles), or Pglyrp1^Cre/wt^ Rosa26^DTA/DTA^ mice (black circles). Individual data are plotted with means ± SD analyzed with two-way ANOVA with Geisser-Greenhouse correction (no sphericity assumed). Where not shown, comparisons were not significant. N = 9 (Rosa26^DTA/DTA^), 11 (Pglyrp1^Cre/wt^ Rosa26^DTA/wt^), and 7 (Pglyrp1^Cre/wt^ Rosa26^DTA/DTA^). Markers used to define each cell population throughout all the graphs are specified in the x-axis. Markers used to identify each cell population are described in Supplemental Table 1B.

Thus, constitutive DTA expression did not deplete *Pglyrp1*-expressing cells in mucosal tissues, suggesting limited toxin efficacy or compensatory survival mechanisms.

### *Pglyrp1*-Cre–driven expression of GFP-DTA results in perinatal lethality and mosaic expression in survivors

Following the lack of detectable cell loss in *Pglyrp1*-Cre–Rosa26^DTA^ mice, we next tested a second DTA-expressing mouse line that also incorporates a Cre-independent GFP reporter upstream of the DTA cassette at the Rosa26 locus. This model, B6.129S6(Cg)-Gt(ROSA)26Sor^tm1(DTA)Jpmb^/J (hereafter Rosa26^GFP-DTA^) [28], enables both functional ablation of Cre-expressing cells and fluorescent labeling of all cells carrying the targeted allele, independent of recombination. Analysis of crosses revealed markedly reduced survival of *Pglyrp1*^Cre/wt^ Rosa26^GFP-DTA/wt^ pups at weaning (Supplemental Figure 5A), with closer examination revealing significant perinatal lethality among *Pglyrp1*^Cre/wt^ Rosa26^GFP-DTA/wt^ pups. Most *Pglyrp1*^Cre/wt^ Rosa26^GFP-DTA/wt^ pups died within the first 24-48 hours of life and deceased pups exhibited abdominal distension with fluid accumulation suggestive of intestinal dysfunction. Marked inflammation and tissue degradation observed in the abdominal region of deceased pups prevented their collection for histological analysis.

Despite repeated breeding attempts, only a small number of double-positive animals survived to weaning. These rare survivors were used to examine the effects of constitutive DTA expression on mucosal cell populations in the gut (Figures 5A, B) and NALT (Figures 5C, D) using the same flow cytometry panel employed in the Tomato reporter experiments (Supplemental Figure 3A). Because M cells are present at low abundance, they were analyzed separately from other epithelial and immune populations in both the gut (Figure 5B) and NALT (Figure 5D). Surprisingly, the analysis revealed no significant differences in overall cell populations (Figure 5A) or M cell subsets (Figure 5B) in the GI tract, nor in corresponding populations in the NALT (Figures 5C, D), when comparing *Pglyrp1*-Cre–positive GFP-DTA mice with control littermates that only contained the Cre-insertion.

**Figure 5.**
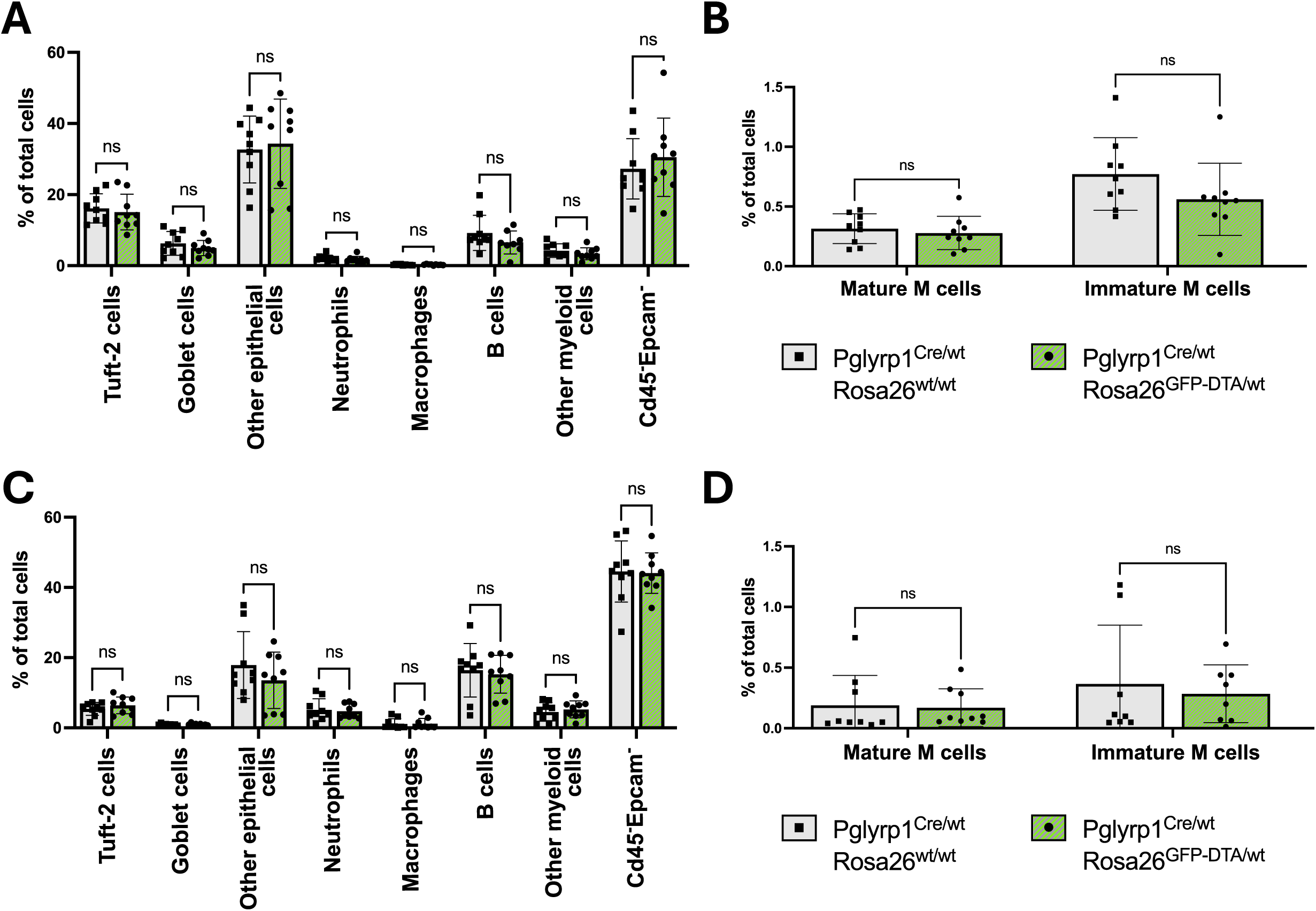
Pglyrp1-Cre–driven expression of GFP-DTA results in perinatal lethality and mosaic expression in survivors. (A) Flow cytometry analysis of cell population levels, and (B) M cell populations in gut tissue of Pglyrp1^Cre/wt^Rosa26^wt/wt^ (gray bars), and Pglyrp1^Cre/wt^Rosa26^GFP-DTA/wt^ (green bars) mice. (C) Flow cytometry analysis of cell population levels, and (D) detailed levels of M cell populations of same mice in NALT. Individual data are plotted with means ± SD analyzed with multiple Mann-Whitney test with Holm-Sidak correction. ns = not significant. N = 9 (Pglyrp1^Cre/wt^Rosa26^wt/wt^), 9 (Pglyrp1^Cre/wt^Rosa26^GFP-DTA/wt^). Markers used to identify each cell population are described in Supplemental Table 1A.

To understand why some double-positive animals survived while others died, we examined GFP expression as a surrogate for DTA expression in GI tract and NALT tissues of surviving mice. Despite being driven by a Cre-independent promoter at the Rosa26 locus, surviving double-positive animals showed mosaic GFP expression, with only 32% of gut cells and 55% of NALT cells expressing GFP, suggesting incomplete transgene activity in surviving mice (Supplemental Figure 5B).

These findings suggest that widespread DTA expression in *Pglyrp1*+ lineages is incompatible with neonatal survival, whereas mosaic expression in survivors limits detectable depletion.

### Assessment of mucosal cell depletion using Rosa26^iDTR^ and diphtheria toxin

To avoid perinatal lethality and allow temporal control of ablation, we used the inducible C57BL/6-Gt(ROSA)26Sor^tm1(HBEGF)Awai^/J strain (hereafter referred to as Rosa26^iDTR^), which expresses the diphtheria toxin receptor (DTR) in a Cre-dependent manner [29]. In this system, *Pglyrp1*-expressing cells remain viable unless selectively depleted by administration of exogenous diphtheria toxin (DT). To determine an appropriate DT dose, we first performed a titration study using intraperitoneal (100, 200, 300, and 500 ng) and intranasal (200, 300, 500, and 1,000 ng) delivery routes (data not shown), following treatment regimens previously reported for the Rosa26^iDTR^ strain [29–32]. At doses above 200 ng, animals exhibited signs of toxicity including perianal bleeding, reduced mobility, and abdominal swelling, in some cases as early as 24 hours after the first administration. Based on these observations, we selected 200 ng as the highest dose that produced no lethality and minimal distress for both intraperitoneal and intranasal administration.

In the gut, intraperitoneal DT increased B cells (Cd45^+^Epcam^-^Ly6g^-^F4/80^-^ B220^+^) regardless of genotype and caused modest, non-significant decreases in some epithelial subsets (Figures 6A, B). M cell populations remained unchanged. In the NALT, DT reduced B cells and neutrophils in *Pglyrp1*^Cre/wt^ Rosa26^iDTR/wt^ mice and increased uncharacterized epithelial cells (Cd45^-^Epcam^+^Gp2^-^Tnfaip2^-^), but again M cell frequencies were unaffected (Figures 6C, D).

**Figure 6.**
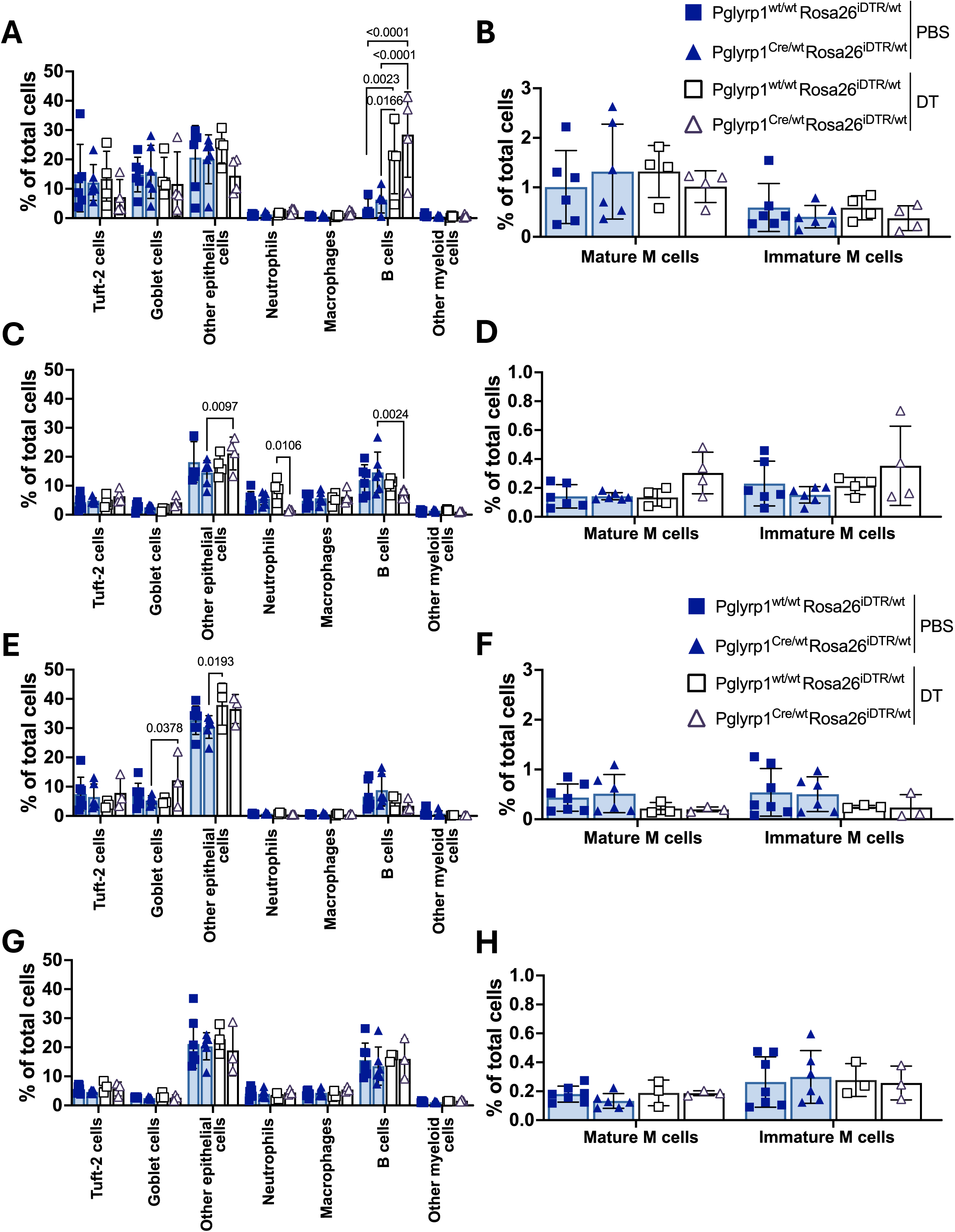
Assessment of mucosal cell depletion using Rosa26^iDTR^ and Pglyrp1-Cre under intraperitoneal and intranasal DT treatment. (A) Flow cytometry analysis of cell population levels, and (B) M cell populations in gut tissue of Pglyrp1^wt/wt^Rosa26^iDTR/wt^ (squares) and Pglyrp1^Cre/wt^Rosa26^iDTR/wt^ (triangles) mice after intraperitoneal inoculation of either PBS or 200 ng DT. (C) Flow cytometry analysis of cell population levels, and (D) M cell populations of same mice in NALT. Individual data are plotted with means ± SD analyzed with two-way ANOVA and Sidak correction for multiple comparisons. Where not shown, comparisons were not significant. N = 6 (PBS), 4 (200 ng DT). (E) Flow cytometry analysis of cell population levels, and (F) M cell populations in gut tissue of Pglyrp1^wt/wt^Rosa26^iDTR/wt^ (squares) and Pglyrp1^Cre/wt^Rosa26^iDTR/wt^ (triangles) mice after intranasal inoculation of either PBS or 200 ng DT. (G) Flow cytometry analysis of cell populations, and (H) M cell populations of same mice in NALT. Individual data are plotted with means ± SD analyzed with two-way ANOVA and Sidak correction for multiple comparisons. Where not shown, comparisons were not significant. N = 6 (PBS), 3 (200 ng DT). Markers used to identify each cell population are described in Supplemental Table 1A.

Intranasal DT produced similarly modest changes in both gut and NALT, and did not significantly alter M cell abundance (Figures 6E–H).

Thus, *Pglyrp1*-Cre–driven DTR expression allows controlled ablation of some populations but fails to deplete M cells or other *Pglyrp1*-expressing lineages at tolerated DT doses.

## Discussion

We generated a *Pglyrp1*-Cre knock-in mouse model designed to provide genetic access to *Pglyrp1*-expressing lineages, including M cells. The knock-in strategy preserved endogenous promoter regulation and resulted in healthy, fertile animals, indicating minimal disruption to *Pglyrp1* function.

Our findings demonstrate that *Pglyrp1* promoter activity is highly tissue specific. In the gut, *Pglyrp1*-Cre marks M cells, goblet cells, and other epithelial subsets, while in the NALT it marks neutrophils and tuft cells more prominently. This underscores the importance of tissue context in interpreting Cre driver activity and highlights the challenge of achieving M cell-specific genetic targeting across mucosal sites.

Functional studies further underscored these limitations. Constitutive DTA expression caused perinatal lethality, suggesting essential roles for *Pglyrp1*-expressing cells early in life, possibly related to gut barrier maturation during neonatal feeding. In contrast, inducible iDTR-mediated ablation resulted in limited depletion at tolerated DT doses, suggesting that incomplete uptake or restricted toxin access constrains effective lineage ablation.

Despite these challenges, *Pglyrp1*-Cre provides a useful tool for probing *Pglyrp1*-expressing populations *in vivo*. Its activity in epithelial and immune compartments enables studies of mucosal homeostasis, innate immune signaling, and barrier function. The lethality observed with GFP-DTA highlights the physiological importance of these lineages during the neonatal period.

More broadly, this work illustrates that M cell-associated gene expression patterns vary across tissues and overlap with other epithelial and immune subsets. Insights from this model will guide future efforts to develop next-generation M cell-specific Cre drivers with improved tissue specificity.

## Material and methods

### Animals

The *Pglyrp1*-Cre mouse was generated using the Easi-CRISPR technique [22], as detailed below, in collaboration with the Childreńs Medical Center Research Institute Genome Engineering Core at UT Southwestern.

The Rosa26^DTA^ strain (B6.129P2-Gt(ROSA)26Sor^tm1(DTA)Lky^/J) [27], Rosa26^GFP-DTA^ strain (B6.129S6(Cg)-Gt(ROSA)26Sor^tm1(DTA)Jpmb^/J) [28], and Rosa26^iDTR^ strain (C57BL/6-Gt(ROSA)26Sor^tm1(HBEGF)Awai^/J) [29] were obtained directly from The Jackson Laboratory. The Tomato reporter strain (B6.Cg-Gt(ROSA)26Sor^tm14(CAG-tdTomato)Hze^/J) [23] was generously provided by Denise Marciano at UT Southwestern. All mice were maintained on a C57Bl/6J genetic background.

Genotyping of mice and embryos was performed according to the protocols provided for each strain by The Jackson Laboratory. The genotyping strategy for the *Pglyrp1*-Cre mouse is described in this manuscript. Briefly, genomic DNA was extracted from tail biopsies of newborn pups or ear punches of weaned mice using a previously described protocol [33].

All experimental procedures were approved by the Institutional Animal Care and Use Committee (IACUC) of UT Southwestern. Mice were housed under specific pathogen-free conditions in accordance with institutional guidelines.

### Generation of *Pglyrp1*-Cre mice

Suitable protospacer adjacent motif (PAM) sites were identified within exon 3 of the *Pglyrp1* locus to guide CRISPR-associated protein 9 (Cas9)-mediated double-strand breaks. The DNA repair template consisted of a 2,015-base pair ssDNA donor containing homology arms flanking the targeted cut site, an IRES, and the coding sequence of Cre recombinase (Figure 1B; Supplemental Figure 1A). The IRES element, provided by the Childreńs Medical Center Research Institute Genome Engineering Core at UT Southwestern, ensured independent translation of Cre recombinase from the endogenous *Pglyrp1* transcript. The Cre sequence (1,026 bp, excluding the start and stop codons) was obtained from the Addgene Sequence Analyzer (https://www.addgene.org/browse/sequence/199783/), and the ssDNA donor construct was synthesized via Genewiz single-stranded DNA synthesis service.

To facilitate genotyping, EcoRI and NheI restriction enzyme recognition sites were inserted at the 5⍰ and 3⍰ ends of the cassette. The endogenous *Pglyrp1* stop codon was preserved, while the Cre sequence was flanked by independent start and stop codons to ensure independent expression of both proteins. Finally, we introduced a silent point mutation within the PAM sequence (highlighted in red, Supplemental Figure 1B) to prevent repeated Cas9 recognition and re-cutting. The required guide RNA was ordered from IDT.

Zygote injections were performed by the Children’s Medical Center Research Institute Genome Engineering Core at UT Southwestern using standard Easi-CRISPR protocols.

### Whole-mount immunofluorescence on newborn pups

The euthanasia and fixation of whole newborn pups were performed in accordance with the guidelines of IACUC of UT Southwestern. Briefly, 1-day-old pups were euthanized, decapitated, and skinned to facilitate the fixation. The entire bodies and heads were fixed with 4% paraformaldehyde (PFA, Thermo Scientific), embedded sagittally in paraffin, sectioned at 5 μm, and mounted on glass slides.

For immunostaining, slides were deparaffinized using xylene and ethanol washes, followed by heat-mediated antigen retrieval in 10mM sodium citrate (pH 6.0). Tissue sections were then permeabilized and blocked for 1 hour at room temperature in 0.4% Triton X-100 and 5% bovine serum albumin (BSA) in PBS (blocking solution with Triton). After washing with PBS, slides were incubated overnight at room temperature with a 1:100 dilution of rabbit anti-RFP (Rockland #D600-401-379S) and rat anti-Gp2 (MBL #D278-3), in blocking solution with Triton. The following day, slides were washed with PBS and incubated for 1 hour at room temperature with a 1:400 dilution of goat anti-rabbit IgG-Alexa Fluor 488 (Invitrogen #A11008) and donkey anti-rat IgG Alexa Fluor 594 (Invitrogen #A21209), in blocking solution with Triton. After a final series of PBS washes, slides were incubated with DAPI, washed again, mounted in Prolong Gold antifade reagent, and imaged using an Axioscan.Z1 slide scanner (Zeiss).

### Flow cytometry

Single-cell suspensions from gut and NALT tissues were prepared using a modified version of a previously described protocol [34]. Briefly, a section of the ileum and the NALT were collected separately, minced with scissors, and digested in RPMI 1640 medium (Gibco)containing 10mM HEPES, 5% heat-inactivated FBS, 30μg/ml Dnase I (Roche), and 1mg/ml Collagenase II (Worthington) at 37°C for 45 minutes. The resulting cell suspension was passed through a 70 μm nylon cell strainer (Falcon #352350), centrifuged, and washed in ACK (Ammonium-Chloride-Potassium) lysis buffer (Gibco #A10492-01). Cells were then resuspended in 5% BSA in PBS (FACSbuffer).

For immunostaining, cells were incubated with the antibody panels specified in each section of the Results, followed by washing and incubation with secondary antibodies in FACS buffer, if needed. After staining, cells were washed and fixed in 4% PFA in PBS for 3 hours (or 1% PFA in PBS overnight) followed by analysis on an LSRII flow cytometer (BD Biosciences) and analyzed using FlowJo software.

The following antibodies were used: rat Epcam (Cd326)-BV421 (1:200, BioLegend #118225), rat Cd45-APC-Cy7 (1:200, Biolegend #103115), rat Ly6g-PE-eFluor610 (1:200, Invitrogen #61-9668-82), rat Ly6G-Alexa Fluor 488 (1:200, BioLegend #127625), rat F4/80-PerCP-Cy5.5 (1:200, Biolegend #123128), rat F4/80-PE (1:200, Biolegend #123110), rat B220-PE-Cy7 (1:200, Biolegend #103221), rabbit Tnfaip2-CF647 (1:100, Biorbyt #orb101923-CF647), rat Cd3-PerCP-Cy5.5 (1:200, BioLegend #155615), rabbit Gp2-FITC (1:100, Biorbyt #orb37776), rabbit Gp2 (1:100, Biorbyt #orb623866), goat anti-rabbit IgG-Alexa Fluor 647 (1:200, Invitrogen #A21245).

### Diphtheria toxin inoculations

Rosa26^iDTR^ mice, with or without *Pglyrp1*-Cre insertion, were injected intraperitoneally for three consecutive days with 100μl of either PBS or 200ng of DT (Sigma) per injection using a 1ml tuberculin syringe (BD #309626).

For intranasal inoculations, the same regimen was followed, with each mouse receiving either 10μl of PBS or 200ng of DT (Sigma) via a 2-20μl pipette.

All mice were 8-11 weeks of age, and all animals were sex- and age-matched. Each experiment was repeated at least twice.

### Statistical analysis

Statistical analyses were performed using GraphPad Prism (RRID:SCR_002798). For flow cytometry studies, two-way ANOVA with correction for multiple comparisons was used when three experimental groups were present. When only two groups were compared, multiple Mann-Whitney tests with Holm-Sidak correction were performed. Survival studies were analyzed using Kaplan-Meier analysis. P-values are indicated in the figures; otherwise, comparisons were not significant. Statistical significance was defined as follows: *p<0.05, **p<0.005, ***p<0.0005, ****p<0.0001.

## Acknowledgements

The authors thank the core facilities at UT Southwestern Medical Center for their valuable contributions to this work, including the Childreńs Medical Center Research Institute Genome Engineering core, particularly to the core director Hao Zhu, and Lin Li, Yu Zhang, and Tripti Sharma; the Histopathology Core, particularly to the core director Bret Evers, and John Shelton; the Whole Brain Microscopy Facility (RRID:SCR_017949), particularly to Denise Ramirez; the Flow Cytometry Facility, particularly to David Farrar; and the Moody Foundation Flow Cytometry Facility for providing access to GraphPad Prism software.

## Author contributions

Conceptualization, S.A.A., M.U.S.; Formal analysis, S.A.A., M.U.S.; Investigation, S.A.A.; Funding acquisition, M.U.S.; Project administration, S.A.A., M.U.S.; Supervision, M.U.S.; Writing – original draft, review and editing, S.A.A., M.U.S.

## References

1. Roe, K., The epithelial cell types and their multi-phased defenses against fungi and other pathogens. Clin Chim Acta, 2024. 563: p. 119889.

2. Kobayashi, N., et al., The Roles of Peyer’s Patches and Microfold Cells in the Gut Immune System: Relevance to Autoimmune Diseases. Front Immunol, 2019. 10: p. 2345.

3. Mabbott, N.A., et al., Microfold (M) cells: important immunosurveillance posts in the intestinal epithelium. Mucosal Immunol, 2013. 6(4): p. 666–77.

4. Corr, S.C., C.C. Gahan, and C. Hill, M-cells: origin, morphology and role in mucosal immunity and microbial pathogenesis. FEMS Immunol Med Microbiol, 2008. 52(1): p. 2–12.

5. Teitelbaum, R., et al., The M cell as a portal of entry to the lung for the bacterial pathogen Mycobacterium tuberculosis. Immunity, 1999. 10(6): p. 641–50.

6. Nair, V.R., et al., Microfold Cells Actively Translocate Mycobacterium tuberculosis to Initiate Infection. Cell Rep, 2016. 16(5): p. 1253–1258.

7. Khan, H.S., et al., Identification of scavenger receptor B1 as the airway microfold cell receptor for Mycobacterium tuberculosis. Elife, 2020. 9.

8. Dillon, A. and D.D. Lo, M Cells: Intelligent Engineering of Mucosal Immune Surveillance. Front Immunol, 2019. 10: p. 1499.

9. Knoop, K.A., et al., RANKL is necessary and sufficient to initiate development of antigen-sampling M cells in the intestinal epithelium. J Immunol, 2009. 183(9): p. 5738–47.

10. Wang, J., et al., Convergent and divergent development among M cell lineages in mouse mucosal epithelium. J Immunol, 2011. 187(10): p. 5277–85.

11. Kanaya, T., et al., The Ets transcription factor Spi-B is essential for the differentiation of intestinal microfold cells. Nat Immunol, 2012. 13(8): p. 729–36.

12. Kimura, S., et al., Sox8 is essential for M cell maturation to accelerate IgA response at the early stage after weaning in mice. J Exp Med, 2019. 216(4): p. 831–846.

13. Madison, B.B., et al., Cis elements of the villin gene control expression in restricted domains of the vertical (crypt) and horizontal (duodenum, cecum) axes of the intestine. J Biol Chem, 2002. 277(36): p. 33275–83.

14. Rios, D., et al., Antigen sampling by intestinal M cells is the principal pathway initiating mucosal IgA production to commensal enteric bacteria. Mucosal Immunol, 2016. 9(4): p. 907–16.

15. Kurashima, Y., et al., Pancreatic glycoprotein 2 is a first line of defense for mucosal protection in intestinal inflammation. Nat Commun, 2021. 12(1): p. 1067.

16. George, J.J., et al., PRC2 Regulated Atoh8 Is a Regulator of Intestinal Microfold Cell (M Cell) Differentiation. Int J Mol Sci, 2021. 22(17).

17. Lo, D., et al., Peptidoglycan recognition protein expression in mouse Peyer’s Patch follicle associated epithelium suggests functional specialization. Cell Immunol, 2003. 224(1): p. 8–16.

18. Haber, A.L., et al., A single-cell survey of the small intestinal epithelium. Nature, 2017. 551(7680): p. 333–339.

19. Dziarski, R. and D. Gupta, Review: Mammalian peptidoglycan recognition proteins (PGRPs) in innate immunity. Innate Immun, 2010. 16(3): p. 168–74.

20. Liu, C., et al., Peptidoglycan recognition proteins: a novel family of four human innate immunity pattern recognition molecules. J Biol Chem, 2001. 276(37): p. 34686–94.

21. Park, S.Y., et al., Peptidoglycan recognition protein 1 enhances experimental asthma by promoting Th2 and Th17 and limiting regulatory T cell and plasmacytoid dendritic cell responses. J Immunol, 2013. 190(7): p. 3480–92.

22. Miura, H., et al., Easi-CRISPR for creating knock-in and conditional knockout mouse models using long ssDNA donors. Nat Protoc, 2018. 13(1): p. 195–215.

23. Madisen, L., et al., A robust and high-throughput Cre reporting and characterization system for the whole mouse brain. Nat Neurosci, 2010. 13(1): p. 133–40.

24. Alvarez-Arguedas, S., et al., Single cell transcriptional analysis of human adenoids identifies molecular features of airway microfold cells. Mucosal Immunol, 2025.

25. Dukhanina, E.A., et al., A new role for PGRP-S (Tag7) in immune defense: lymphocyte migration is induced by a chemoattractant complex of Tag7 with Mts1. Cell Cycle, 2015. 14(22): p. 3635–43.

26. Kang, D., et al., A peptidoglycan recognition protein in innate immunity conserved from insects to humans. Proc Natl Acad Sci U S A, 1998. 95(17): p. 10078–82.

27. Voehringer, D., H.E. Liang, and R.M. Locksley, Homeostasis and effector function of lymphopenia-induced “memory-like” T cells in constitutively T cell-depleted mice. J Immunol, 2008. 180(7): p. 4742–53.

28. Ivanova, A., et al., *In vivo* genetic ablation by Cre-mediated expression of diphtheria toxin fragment A. Genesis, 2005. 43(3): p. 129–35.

29. Buch, T., et al., A Cre-inducible diphtheria toxin receptor mediates cell lineage ablation after toxin administration. Nat Methods, 2005. 2(6): p. 419–26.

30. Bruttger, J., et al., Genetic Cell Ablation Reveals Clusters of Local Self-Renewing Microglia in the Mammalian Central Nervous System. Immunity, 2015. 43(1): p. 92–106.

31. Rodriguez-Baena, F.J., et al., Microglial reprogramming enhances antitumor immunity and immunotherapy response in melanoma brain metastases. Cancer Cell, 2025.

32. Gritsch, S., et al., Oligodendrocyte ablation triggers central pain independently of innate or adaptive immune responses in mice. Nat Commun, 2014. 5: p. 5472.

33. Truett, G.E., et al., Preparation of PCR-quality mouse genomic DNA with hot sodium hydroxide and tris (HotSHOT). Biotechniques, 2000. 29(1): p. 52, 54.

34. Wolf, A.J., et al., Mycobacterium tuberculosis infects dendritic cells with high frequency and impairs their function *in vivo*. J Immunol, 2007. 179(4): p. 2509–19.

