## Supplemental Figures for "Pglyrp1-Cre Marks Distinct Epithelial and Immune Lineages Across Mucosal Sites"

### Supplemental Figure 1

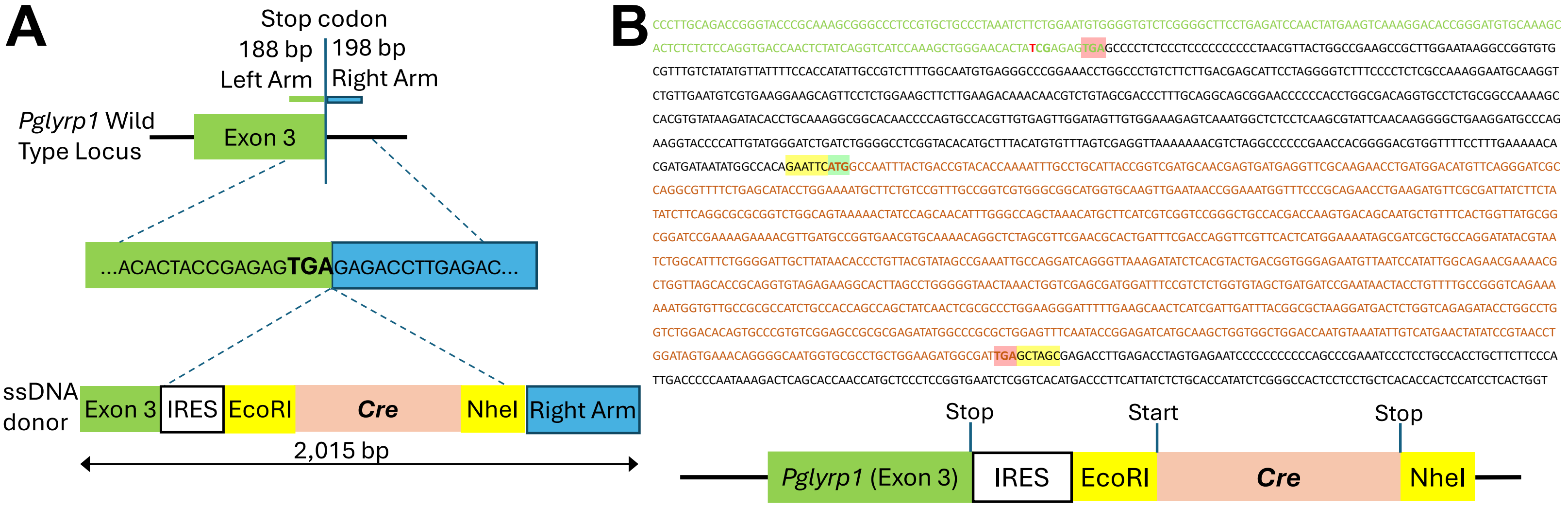


**C**

| | 5´ Fw – 3´ Rv | 5´ Fw – 5´ Rv | 3´ Fw – 3´ Rv |
|---|---|---|---|
| WT | 170 bp | -- | -- |
| KI | -- | 733 bp | 326 bp |
| Enz. digestion | 170 bp | 655 bp + 78 bp | 220 bp + 106 bp |

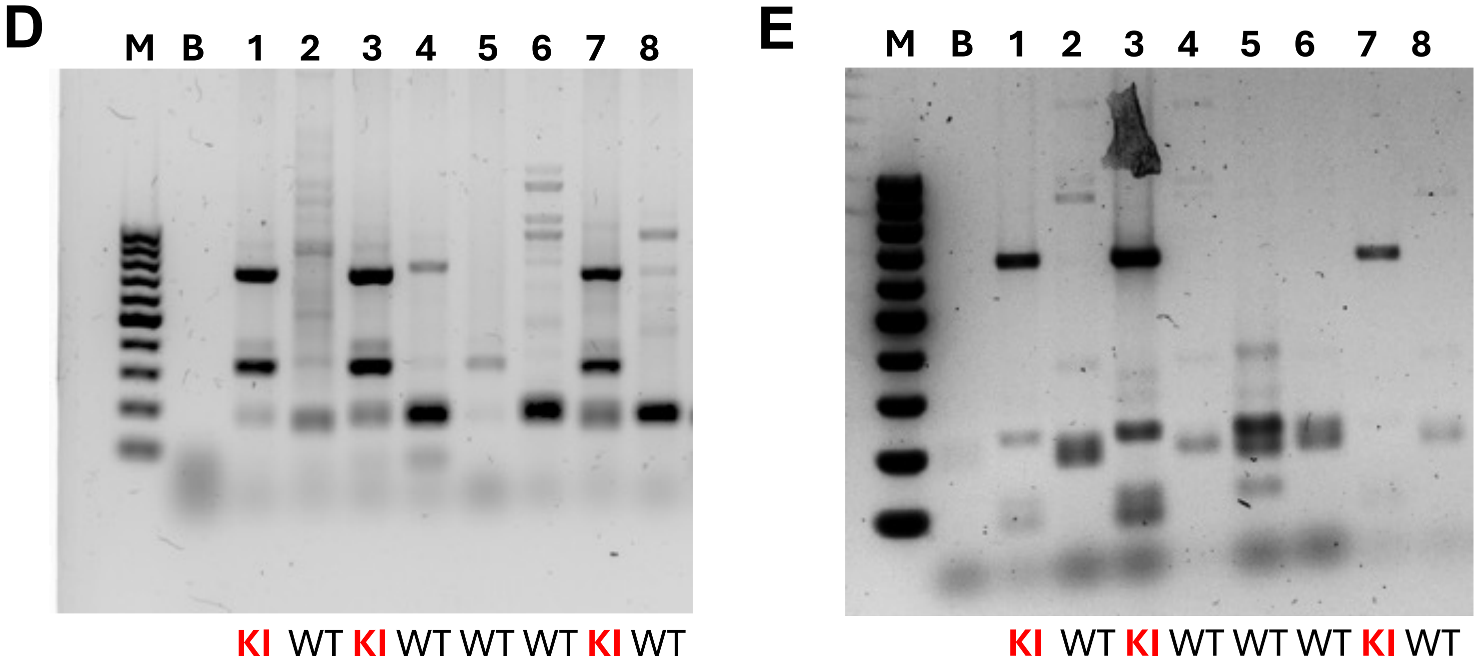

### Supplemental Figure 2

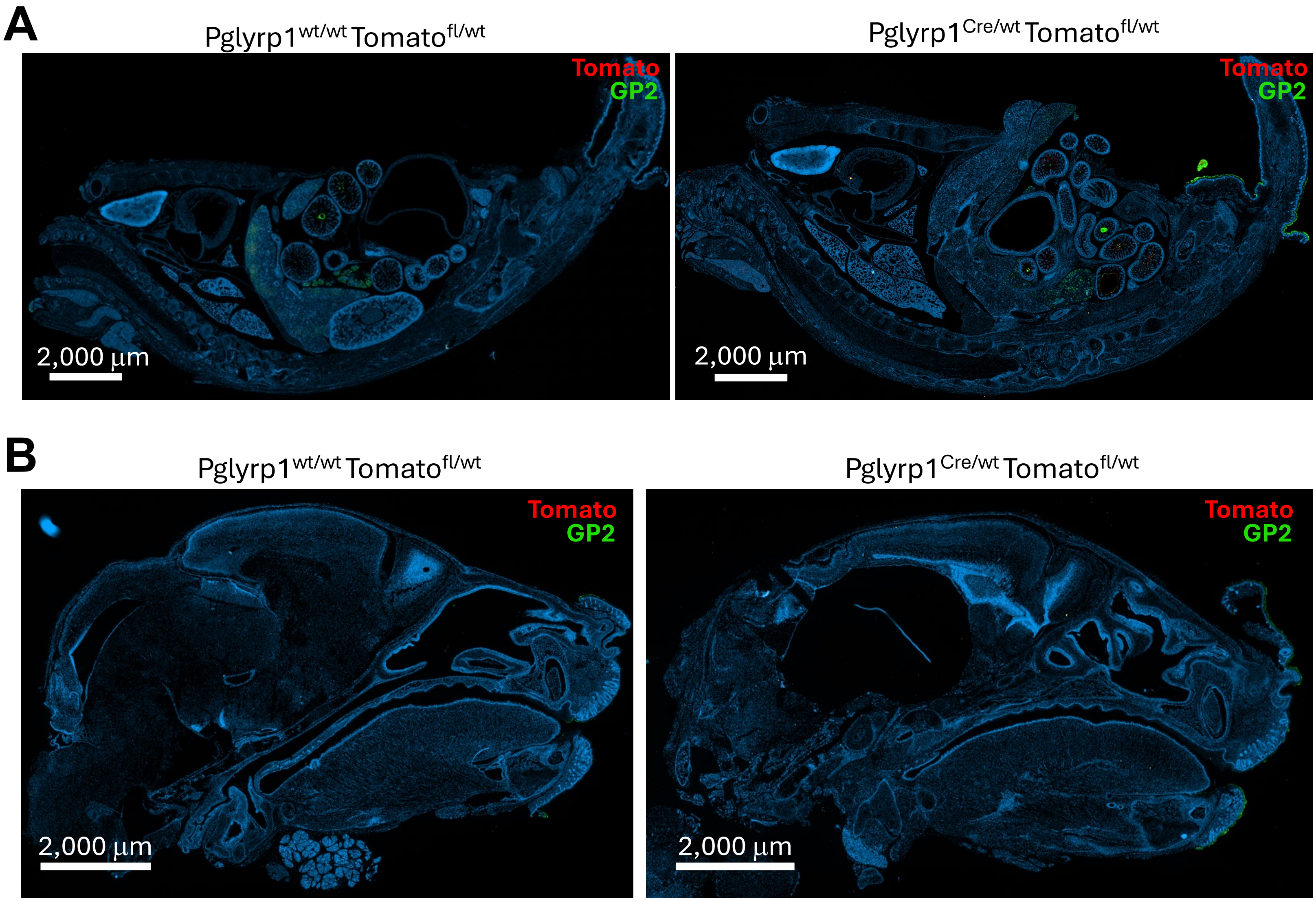

### Supplemental Figure 3

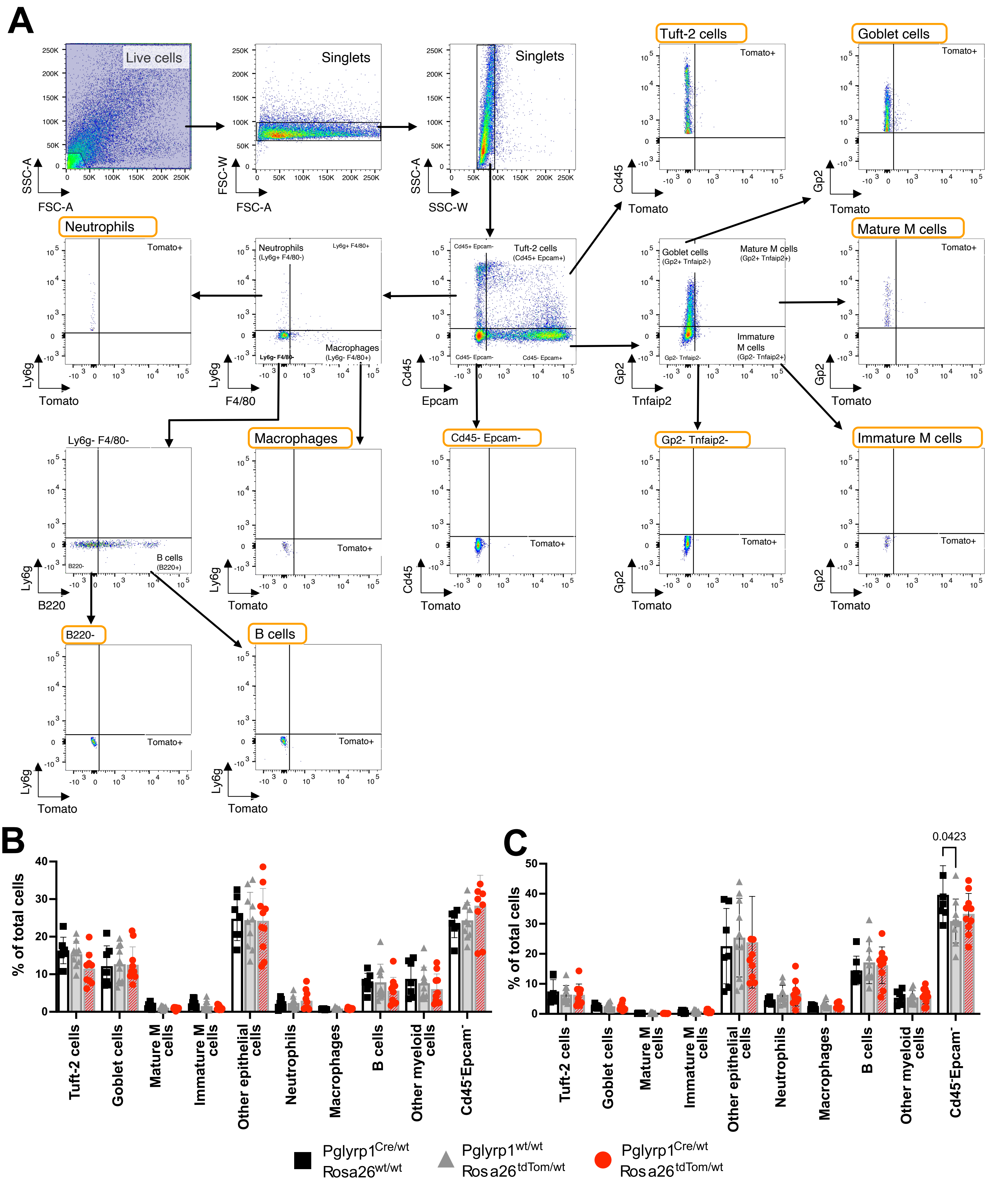

### Supplemental Figure 4

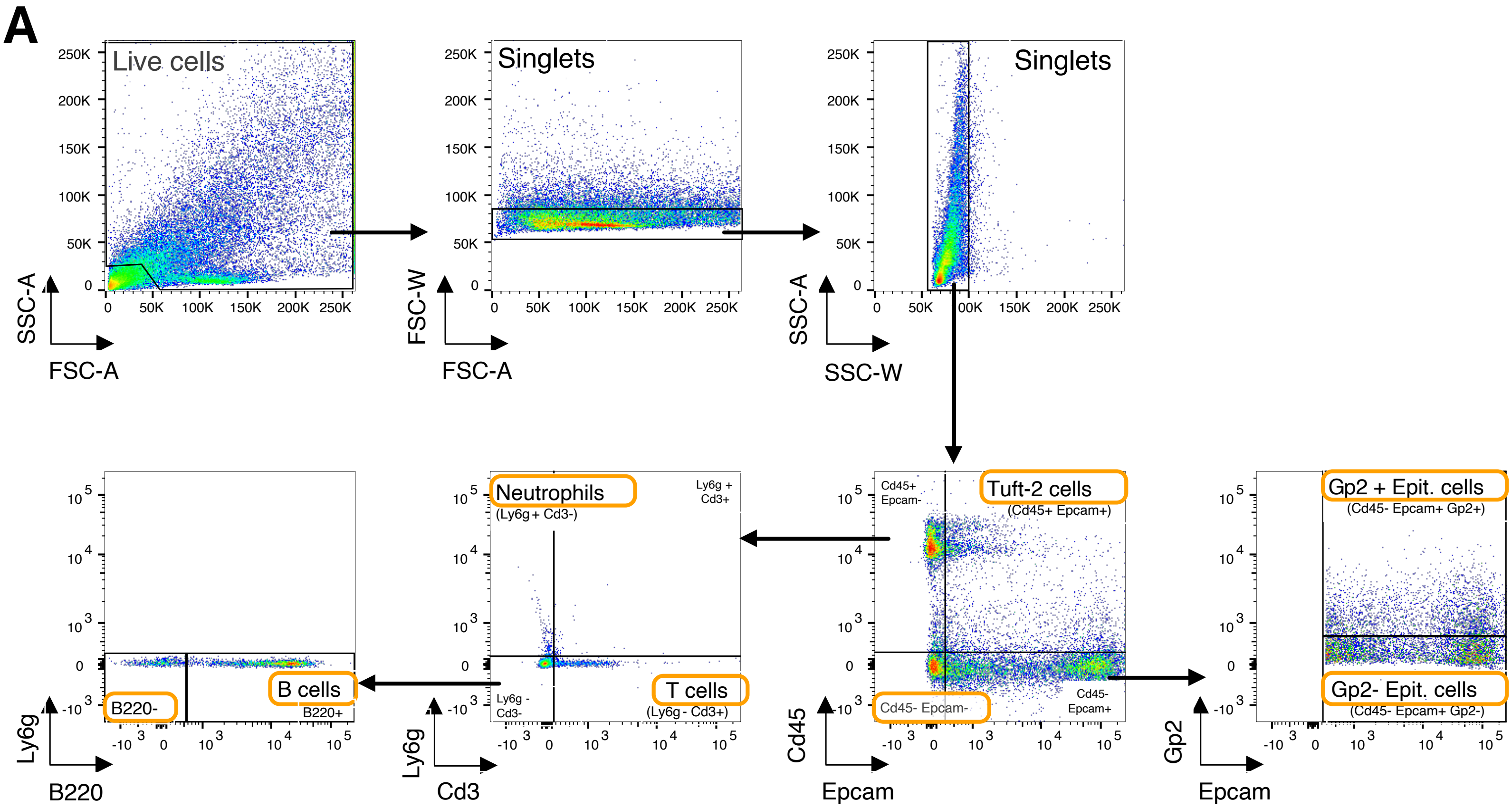

### Supplemental Figure 5

**A**

| $F_0$ | $F_1$ probability | #pups at weaning | $F_1$ genotypes |
|---|---|---|---|
| $Rosa26^{GFP\text{-}DTA/wt}$ x $Pglyrp1^{Cre/wt}$ | **25%** $Pglyrp1^{Cre/wt}$ $Rosa26^{GFP\text{-}DTA/wt}$ | 24 | 0 |
| | **25%** $Pglyrp1^{Cre/wt}$ $Rosa26^{wt/wt}$ | | 13 |
| | **25%** $Pglyrp1^{wt/wt}$ $Rosa26^{GFP\text{-}DTA/wt}$ | | 4 |
| | **25%** $Pglyrp1^{wt/wt}$ $Rosa26^{wt/wt}$ | | 7 |

**B**

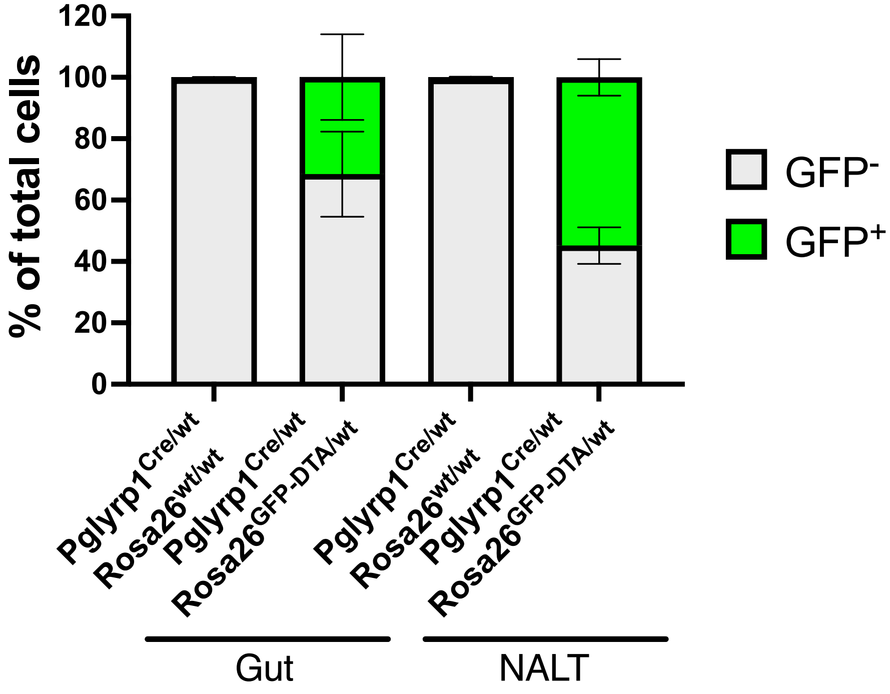

### Supplemental Table 1

## A

**Markers for flow cytometry in figures 3, 5, and 6.**

| Cell population | Identifying biomarkers |
|---|---|
| Tuft cells | $Cd45^{+}$ $Epcam^{+}$ |
| Goblet cells | $Cd45^{-}$ $Epcam^{+}$ $Gp2^{+}$ $Tnfaip2^{-}$ |
| Mature M cells | $Cd45^{-}$ $Epcam^{+}$ $Gp2^{+}$ $Tnfaip2^{+}$ |
| Immature M cells | $Cd45^{-}$ $Epcam^{+}$ $Gp2^{-}$ $Tnfaip2^{+}$ |
| Other epithelial cells | $Cd45^{-}$ $Epcam^{+}$ $Gp2^{-}$ $Tnfaip2^{-}$ |
| Neutrophils | $Cd45^{+}$ $Epcam^{-}$ $Ly6g^{+}$ |
| Macrophages | $Cd45^{+}$ $Epcam^{-}$ $Ly6g^{-}$ $F4/80^{+}$ |
| B cells | $Cd45^{+}$ $Epcam^{-}$ $Ly6g^{-}$ $F4/80^{-}$ $B220^{+}$ |
| Other myeloid cells | $Cd45^{+}$ $Epcam^{-}$ $Ly6g^{-}$ $F4/80^{-}$ $B220^{-}$ |
| | $Cd45^{-}$ $Epcam^{-}$ |

## B

**Markers for flow cytometry in figure 4.**

| Cell population | Identifying biomarkers |
|---|---|
| Tuft cells | $Cd45^{+}$ $Epcam^{+}$ |
| $Gp2^{+}$ epithelial cells | $Cd45^{-}$ $Epcam^{+}$ $Gp2^{+}$ |
| $Gp2^{-}$ epithelial cells | $Cd45^{-}$ $Epcam^{+}$ $Gp2^{-}$ |
| Neutrophils | $Cd45^{+}$ $Epcam^{-}$ $Ly6g^{+}$ |
| T cells | $Cd45^{+}$ $Epcam^{-}$ $Ly6g^{-}$ $CD3^{+}$ |
| B cells | $Cd45^{+}$ $Epcam^{-}$ $Ly6g^{-}$ $CD3^{-}$ $B220^{+}$ |
| Other myeloid cells | $Cd45^{+}$ $Epcam^{-}$ $Ly6g^{-}$ $CD3^{-}$ $B220^{-}$ |
| | $Cd45^{-}$ $Epcam^{-}$ |

**Supplemental Figure 1. Generation of a *Pglyrp1*-Cre mouse model and founder screening. Related to figure 1.** (A) Genetic sequence of *Pglyrp1* exon 3 in wild-type animals and after repair with the Cre-containing ssDNA donor sequence. (B) Detailed DNA sequence of the 2,015bp ssDNA donor, including left and right homology arms. Color coding corresponds to the schematic representation of the ssDNA donor construct: stop codons are highlighted in red, the *Cre* start codon in green, and restriction enzymes sites –EcoRI and NheI– in yellow. A silent point mutation (thymine, highlighted in red) was introduced to eliminate the original PAM site preventing repeated Cas9 recognition. (C) Table of expected fragment sizes (in base pairs) for each genotype, based on amplification with designed primers and subsequent enzymatic digestion. (D) PCR amplification products obtained from genomic DNA for wild-type and knock-in genotypes, and (E) after enzymatic digestion of the amplified fragments with genotype interpretation provided below each gel image. Lanes containing Cre insertions are highlighted in red. IRES = Internal ribosome entry site, M = 100 bp marker, B = blank, WT = wild type, KI = knock-in.

**Supplemental Figure 2. Reporter expression in newborn *Pglyrp1*-Cre mice reveals tissue-specific patterns. Related to figure 2.** Confocal images of representative Pglyrp1$^{wt/wt}$Rosa26$^{tdTom/wt}$ (left), and Pglyrp1$^{Cre/wt}$Rosa26$^{tdTom/wt}$ (right) one-day-old pups from the whole body (A) and head (B) sections stained with anti-red fluorescent protein (RFP), and anti-GP2 antibodies. Scale bars included in the pictures.

**Supplemental Figure 3. Quantitative analysis of *Pglyrp1*-Cre driven Tomato expression in adult mice highlights tissue-specific patterns. Related to figure 3.** (A) Gating strategy for the detection of Tomato reporter expression in mice in both GI tract and NALT. The figure shows a representative dot plot from GI tract tissue of a Pglyrp1$^{Cre/wt}$Rosa26$^{wt/wt}$ mouse. Main cell populations are highlighted with orange boxes. Percentage of total cells in gut (B), and NALT (C), in Pglyrp1$^{Cre/wt}$ (black squares), Rosa26$^{tdTom/wt}$ (gray triangles), and Pglyrp1$^{Cre/wt}$Rosa26$^{tdTom/wt}$ (red circles) adult mice. Markers used to define each cell population are specified in the x-axis. Individual data are plotted with means ± SD analyzed with two-way ANOVA with Geisser-Greenhouse correction (no sphericity assumed). Where not shown, comparisons were not significant. N = 7 (Pglyrp1$^{Cre/wt}$) and 10 (Rosa26$^{tdTom/wt}$,

Pglyrp1$^{Cre/wt}$Rosa$^{tdTom/wt}$). Markers used to identify each cell population are described in Supplemental Table 1A.

**Supplemental Figure 4. Pglyrp1-Cre–Rosa26$^{DTA}$ mice show no detectable changes in epithelial or immune cell composition. Related to Figure 4.** (A) Gating strategy used for flow cytometric analysis of epithelial and immune cells from the GI tract and NALT. The figure shows a representative dot plot from gut tissue of a Rosa26$^{DTA/DTA}$ mouse. Main cell populations are highlighted with orange boxes.

**Supplemental Figure 5. Pglyrp1-Cre–driven expression of GFP-DTA results in perinatal lethality and mosaic expression in survivors. Related to Figure 5.** (A) Mendelian genotype proportions and number of offspring obtained at weaning (4 weeks after birth) from crosses between *Pglyrp1*-Cre and Rosa26$^{GFP\text{-}DTA}$ mice. (B) Flow cytometry analysis of the percentage of GFP-positive cells in gut and NALT tissues from Pglyrp1$^{Cre/wt}$ animals with or without Rosa26$^{GFP\text{-}DTA}$ allele. N = 9 per group.

**Supplemental Table 1. Markers used to identify each cell population in flow cytometry.** (A) List of cell populations and their corresponding identifying biomarkers used in figures 3, 5, and 6. (B) List of cell populations and their corresponding identifying biomarkers used in figure 4.
